# Pervasive integrative and conjugative elements shape *Porphyromonas gingivalis* gene repertoires

**DOI:** 10.64898/2026.08.04.741601

**Authors:** Cole B. Matrishin, Elaine M. Haase, Ashley K. Miles, Stephanie Steimer, Dam Soh, Matthew Smardz, Patricia I. Diaz, Kathryn M. Kauffman

## Abstract

**Background:** *Porphyromonas gingivalis* (*Pg*) is an oral pathobiont that contributes to periodontal disease and has been associated with systemic health conditions. Although *Pg* is recognized as exhibiting extensive strain-level genomic diversity and recombination, the extent to which mobile elements contribute to this variation, and their relevance to its fitness and virulence, remain incompletely understood. Our recent study of the *Pg* pangenome revealed diverse accessory defense-associated genes, raising the question of whether these are carried by unrecognized mobile genetic elements (MGEs). Integrative and conjugative elements (ICEs) are large autonomous mobile elements that often encode genes for proteins beneficial to their bacterial hosts, including defense systems that protect against phage infection. To date, only one ICE, CTnPg1, has been described in *Pg*.

**Results:** Here, we developed a bioinformatic approach integrating ICE prediction and curation, hallmark-gene detection, and genomic-context analysis, to investigate ICEs in *Pg*. We discovered that ICEs are pervasive in *Pg* genomes, with >90% of genomes harboring at least one ICE. We found that these elements comprise at least five distinct groups, two of which dominate and frequently co-occur in *Pg* genomes, inserting into distinct characteristic insertion sites. Using marker-gene analysis of enrichment-culture mini-metagenomes from subjects with periodontal disease we detected representatives of these dominant *Pg* ICE groups, as well as others, in recent clinical samples. We found that anti-defense and defense genes are common in *Pg* ICEs, and that these elements commonly encode biosynthetic gene clusters, including for menaquinone synthesis and predicted ribosomally synthesized and post-translationally modified peptides (RiPPs). In contrast to the extensive CRISPR-Cas defense targeting we observed for *Pg* phages, we detected no exact matches between ICE sequences and *Pg* CRISPR spacers.

**Conclusion:** This work establishes that ICEs are pervasive contributors to *Pg*’s pangenome and unique strain-level gene repertoires. Their distinct cargo profiles suggest that ICEs likely impact the virulence and ecology of *Pg* through the introduction and spread of advantageous traits, including expansion of *Pg*’s biosynthetic capacity and resistance to phage infection. This work provides a curated framework for investigating ICE diversity in *Pg* and establishes a foundation for expanded experimental studies of their host ranges and roles in shaping *Pg*’s interactions with phages, other microbes, and the human host.

## INTRODUCTION

The human oral microbiome is one of the most well-studied and characterized microbial communities^1^. Complex interactions of its over 600 bacterial species, within and across the distinct ecological niches of the mouth^2^, have been intensively studied and provide a rich context for linking microbial community structure and function with human health. Shifts in the microbial composition of these communities, known as dysbiosis, can result in oral disease, including periodontal disease, a chronic inflammatory condition that results in destruction of the tooth-supporting soft tissues and bone, with its severe form affecting ∼10% adults 30 or older in the United States^3^.

*Porphyromonas gingivalis* (*Pg*) is a Gram-negative, anaerobic pathogen that acts as a keystone driver of periodontal disease^4,5^. *Pg* colonizes subgingival polymicrobial biofilms, relying on coaggregation with other bacteria and uptake of growth factors produced by neighbors^6^. Once established in the subgingival crevice, *Pg* can create a pro-inflammatory environment through the release of cysteine proteases, known as gingipains, that facilitate nutrient acquisition, immune evasion, and tissue destruction^4,5^. *Pg* has also been associated with many systemic conditions^7^, including cardiovascular disease^8^, adverse pregnancy outcomes^9,10^, and Alzheimer’s disease^11,12^, highlighting the importance of understanding the factors that allow *Pg* to colonize and persist within the oral microbiome, and disseminate systemically. However, observations of *Pg* in healthy donors, and evident genomic and phenotypic variation between isolates^13^, underscore the need for a better understanding of the factors that contribute to *Pg* strain-level diversity.

Our recent study of the *Pg* pangenome, as part of our investigation of its infecting prophages, found that nearly a fourth of every *Pg* strain’s genome is composed of ‘flexible’ or ‘accessory’ genes not shared by all members of the species^14^. We identified prophages as contributors to this flexible gene pool, identifying and characterizing the first three families of phages described for *Pg,* as well as showing that they are heavily targeted by Pg’s CRISPR-Cas defense system spacers. In addition, we found that the *Pg* pangenome includes diverse defense system genes beyond CRISPR-Cas. The known association of mobile elements with diverse defense systems^15^, together with our challenges in reliably cultivating phages despite demonstrating they are active in culture, raised the question of whether the presence of other cryptic integrated mobile elements in *Pg* might be shaping this species’ interactions with its phages.

Mobile genetic elements (MGEs) shape bacterial strain-level diversity and contribute to rapid bacterial adaptation^16^. These selfish elements^15^, including phages, plasmids, integrative and conjugative elements, and some transposons and integrons, can alter their bacterial host’s physiology, virulence potential, and interactions with other microbes and the human host. These effects arise both from expansion of their host’s genetic repertoire through MGE-encoded functions, and from disruption of genes or their regulation during integration^17^. However, uncovering the impact of MGEs on bacterial strains and species can be a challenge, in part due to technical difficulties in identifying and functionally characterizing their sequences in bacterial genomes.

In *Pg*, the three major groups of MGEs that have been described include insertion sequence (IS) elements, phages^14^, and integrative conjugative elements (ICEs)^18,19^, the latter dubbed conjugative transposons. IS elements have been shown to play important roles in promoting survival of bacterial populations *in vivo* during phage predation^20,21^, however confer this through gene disruption rather than encoding defense systems. By contrast, ICEs are large autonomous mobile elements that typically carry cargo genes that provide a selective advantage to their host cells, including, for example, resistance to antibiotics and heavy metals, alternate metabolic pathways, virulence determinants, and protection against infection by phages and other MGEs, thus, promoting their own maintenance within the host genome^22^. ICEs integrate into bacterial chromosomes and facilitate their own transfer between bacterial cells via a self-encoded conjugation module, consisting of a relaxosome mobilization complex (MOB) that mobilizes the element and a type IV secretion system (T4SS, or Mating Pair Formation, MPF, system) that transfers the entire element genome to a recipient cell through a physical structure that spans cells^23^. Notably, a 2022 study by Johnson et al.^24^ demonstrated that an anti-phage gene expressed by an ICE in a *Bacillus subtilis* model system conferred protection against killing by phages, both during lytic infection and in lysogenic induction of prophages; *B. subtilis* strains lacking the ICE were susceptible to killing by phages.

To date, only a single ICE, CTnPg1, has been characterized in *Pg*^18,25^. For clarity, we note that “CTn” (a reference to “conjugative transposon”) is a misnomer when applied to the described *Pg* ICE, because a site-specific integrase, rather than a transposase, mediates its integration. Found in the genome of *Pg* strain ATCC 33277, CTnPg1 has been shown to transfer from ATCC 33277 to W83, as well as to other species, such as *Bacteroides thetaiotaomicron* and *Prevotella oralis*, and has recently been shown to exist within oral microbial communities even in the absence of detectable *Pg*^19,26^. CTnPg1-like ICEs were also found in other oral *Bacteroidales*, including *Porphyromonas endodontalis*, *Prevotella buccae*, and *Prevotella intermedia*, emphasizing their active role in horizontal gene transfer in the mouth^19^. An earlier study^27^ identified a *tra* locus encoding the T4SS in W83, however a homologous locus was not present in ATCC 33277. Despite this, conjugal transfer of DNA bidirectionally between the two strains was still successful. This observation, coupled with the characterization of CTnPg1 in ATCC 33277, suggests that diverse ICEs are present in *Pg* genomes and may be contributing to genetic exchange and bacterial adaptation, including resistance to phage infection, in *Pg*^27^.

The goal of the present study was to investigate whether ICEs are more diverse and widespread in *Pg* than currently recognized and, if so, to characterize the nature of their genetic cargo. Using a comprehensive and integrative bioinformatic approach, we determined that ICEs are prevalent across diverse *Pg* genomes and represent at least five distinct groups - with the two most prevalent showing distinct insertion patterns, gene cargos, and predicted host ranges. In contrast to the extensive targeting of phages, these ICEs do not appear to be targeted by *Pg*’s CRISPR-Cas systems. We found that ICEs encode anti-defense proteins that may facilitate their own spread, as well as defense systems that may protect their hosts against phages and other competing MGEs. We further found that the cargo of *Pg*’s ICEs also includes key metabolic and biosynthetic genes predicted to positively impact *Pg*’s fitness. Altogether, we establish that ICEs are substantial contributors to *Pg*’s strain-level diversity, with the potential to shape Pg’s fitness and interactions in the oral microbiome through the acquisition and dissemination of adaptive traits.

## RESULTS

### The majority of sequenced Pg genomes harbor ICE mobile elements

To address the question of how prevalent ICEs are in *Pg* genomes, we sought to systematically evaluate all genomes of the species in the expanded Human Oral Microbiome Database^28,29^ (*e*HOMD v11.02) for these mobile elements. However, as for most species, the preponderance of available genomes comprise numerous contigs rather than single closed assemblies. As MGEs, including ICEs, are often disproportionately impacted by fragmentation - and thereby split across multiple contigs - it was necessary to ‘scaffold’ contigs to determine their sequential order and thereby enable identification and characterization of full length ICEs that span fragments. Yet, an additional challenge in highly recombining species, such as *Pg*, is identifying the optimal reference sequence to scaffold contigs against.

To overcome the challenges of scaffolding, and enable our baseline discovery survey for ICEs, we developed a bioinformatic strategy that leveraged the diversity of the available *Pg* genome collection. In exploratory studies we found that attempts to use nearest neighbors as references failed to efficiently recover ICE regions. We therefore adopted an agnostic all-by-all approach, wherein each of the 63 multi-contig Pg assemblies were scaffolded against all ∼98 non-self and non-identical Pg assemblies, yielding a total of 6,079 scaffolded assemblies for evaluation. Next, to identify the single best scaffolded assembly for each genome, we performed preliminary ICE conjugation system predictions using CONJscan^30^ to aid in selecting those scaffolded assemblies with the least number of broken ICEs for each genome. All scaffolded assemblies were then evaluated using a tiered ranking system that prioritized, for each genome, the scaffold version in which the greatest number of complete conjugative systems were detected (i.e. with the most ICEs pieced back together) and for which the overall quality of scaffolding was highest (see “Materials and Methods”). The single best-scaffolded assembly for each *Pg* genome, from our large-scale screen, was then selected for the final analyses of ICE regions. Each contig from each best-scaffolded assembly was manually submitted to ICEfinder 2.0 on the ICEberg 3.0^31^ webserver for ICE detection, yielding our core set of region ‘hits’ (with additional boundary curation described further below).

Using this systematic approach we found that *Pg* genomes nearly universally harbor one or more ICEs. We found that 91% of *Pg* genomes (72/79) harbor at least one ICE, and 63% (50/79) encode more than one ICE, and in some cases up to four (Fig. 1). We note that in evaluating percentages, and for downstream analyses, we defined a subset of non-redundant isolate genomes (*Pg*_set_79) from the total set of all *Pg* genomes available on *e*HOMD (*Pg*_set_99, Supplementary Data File 1), that excludes re-sequenced strains, derivative sub-strains, and metagenome-assembled genomes (MAGs). We noted that MAGs are depleted in ICEs, likely reflecting loss of fragmented MGE contigs in final MAG bins for each strain.

**Figure 1.**
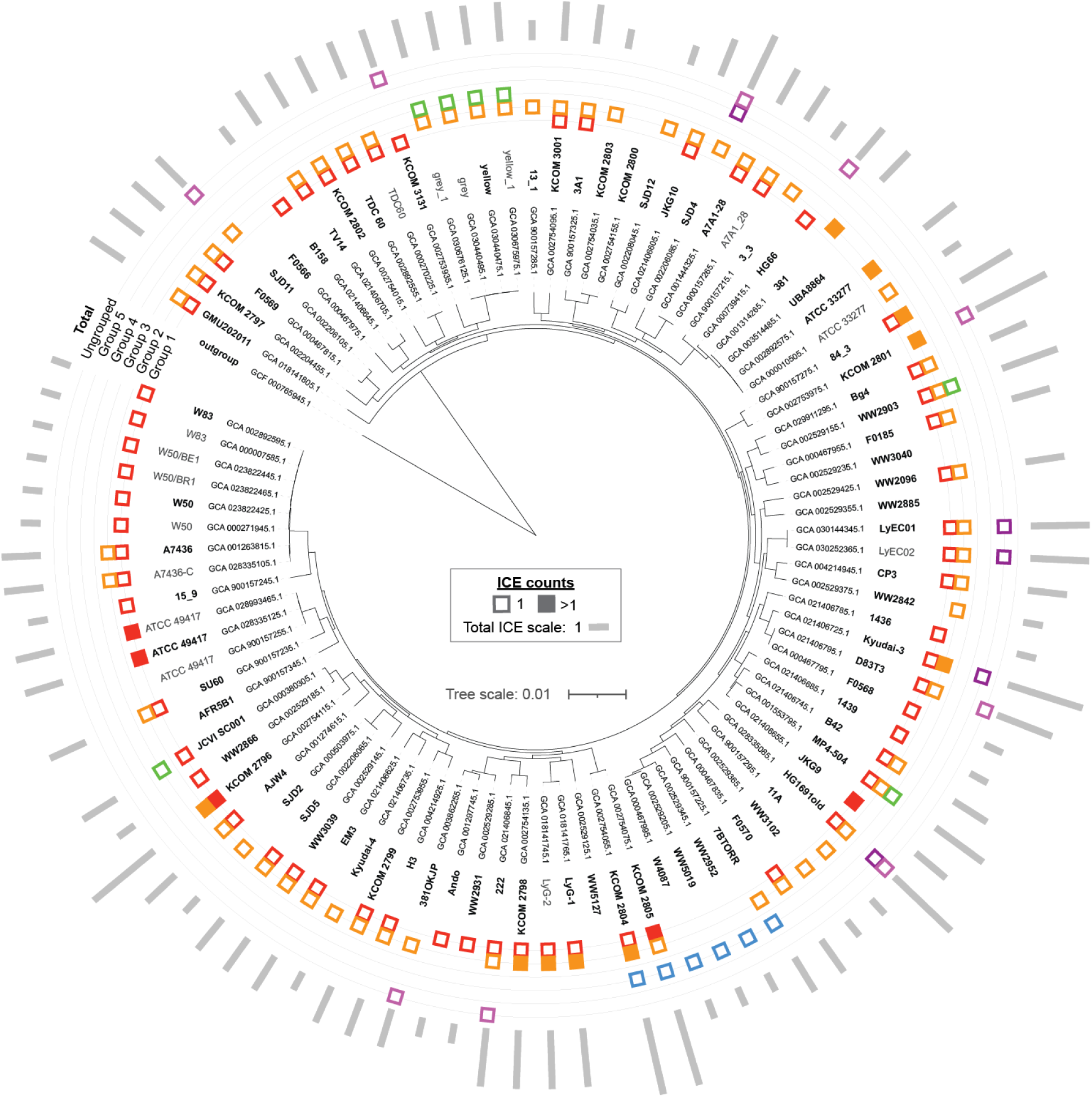
ICEs are widespread and abundant in *Pg* genomes. Phylogenetic tree of *Pg* generated by Panaroo^32^ based on core gene nucleotide alignment. Strains in *Pg*_set_99 are represented (excluding five metagenomic assemblies), with non-redundant strains in *Pg*_set_79 bolded. Inner rings denote the presence of ICEs identified within each strain, with colored boxes respective to the ICE group (classified by 70% nucleotide similarity of conjugation modules determined by VIRIDIC^33^). Empty boxes represent a single ICE, while filled boxes represent multiple ICEs. Outer ring represents the total number of ICEs encoded in each strain. *P. gulae* COT-052 OH3856 (GCF_00765945.1) was used as the outgroup. An interactive version of the iTOL^34^ tree can be found online at https://itol.embl.de/tree/70177211213255061785532670.

Next, we probed the diversity of ICEs, focusing on conserved conjugation module genes, and we found that they separate into five distinct groups (Fig. 2, Supplementary Data File 1). To define operational taxonomic units (OTUs), we used approaches similar to other studies in classifying ICEs and defining OTUs^35–37^. We evaluated conjugation modules, gene regions encompassing the relaxosome [MOB] and mating pair formation [MPF] system genes, including all *mob* and *tra* (T4SS) genes and spanning the region from *mobC* to *traQ*. With this approach, we found that *Pg* ICEs comprise at least five “genus-level” groups (hereafter, simply “groups”) and 47 “species-level” groups, when applying the respective ≥70% and ≥95% nucleotide identity thresholds adapted from the phage taxonomy framework^38^. The two proteins with the greatest degree of sequence conservation across all groups are MobC, an auxiliary transfer protein contributing to relaxosome assembly^39^, and TraG, an inner membrane protein of the T4SS that contributes to ICE transfer efficiency^40^.

**Figure 2.**
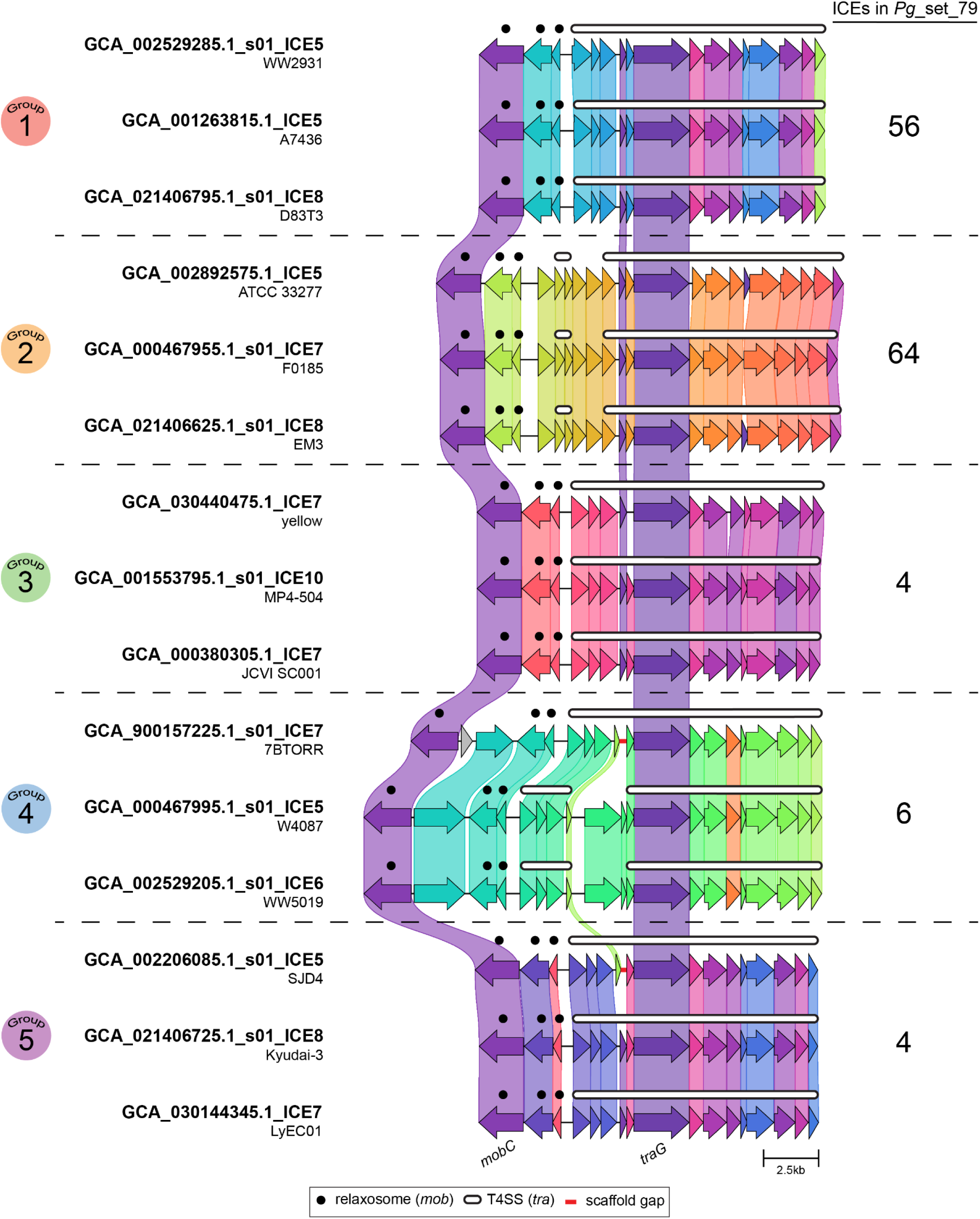
*Pg* ICEs comprise five distinct groups, reflecting distinct conjugation module genes. A subset of conjugation module genes is shown for representative ICEs from each group, with genes colored based on protein sequence similarity and clustering generated by Clinker^41^ (classified by 70% nucleotide identity of conjugation module genes, as determined by VIRIDIC^33^). Colored linkages between genes indicate ≥69% protein sequence similarity. Relaxosome components denoted by black dots and T4SS components denoted by white lines. The total number of ICEs identified within *Pg* genomes from *Pg*_set_79 is shown for each group (nine ungrouped ICEs were identified but not shown).

### Two groups of ICEs with distinct features dominate in Pg

Once classified, two groups of ICEs emerge as prominent and highly prevalent in *Pg* (see group level distribution metadata in Fig. 1). Collectively the 79 nonredundant *Pg* genomes harbor 56 Group 1 ICEs and 64 Group 2 ICEs. Group 1 and Group 2 ICEs also frequently co-occur in *Pg* genomes (37/79 genomes harbor at least one of each group). Consistent with this observation, these ICE groups exhibit distinct integration preferences for conserved insertion site motifs (Supplementary Data File 1). Based on precise manual curation of full-length ICE region boundaries, we find that Group 1 ICEs integrate into specific tRNA genes, tRNA-Asp(gtc) and tRNA-Ser(tga), whereas Group 2 ICEs integrate at sites other than tRNAs. The less prevalent ICE groups also integrate into various tRNA genes, Group 3 into tRNA-Ser(tga), Group 4 into tRNA-Asp(gtc), and Group 5 into tRNA-Phe(gaa). Of interest, our recent study of *Pg* prophages revealed that two families of these phages also insert into tRNA genes, with the rare and often decayed ludisviruses inserting into tRNA-Ser and the far more prevalent nixviruses inserting into tRNA-Pro. These observations highlight that, as shown for *E. coli*^42^, diverse mobile elements may share common insertion sites in their host bacterial genomes.

Given the extensive targeting of integrative phages by CRISPR-Cas defense systems in *Pg*, as we recently showed^14^, we wondered whether *Pg* likewise target ICEs as another class of integrative mobile elements. To determine the extent to which ICEs are targeted, we identified the CRISPR spacers within *Pg* genomes with CRISPRCasTyper^43^ (CCTyper) and mapped them back to the fully-curated ICEs defined in this study (see “Materials and Methods”). By contrast to the extensive targeting of phages, we found that none of the identified spacers (6,320) within the *Pg*_set_99 sequences mapped back to any ICE in our curated set (*Pg*_ICE_pilot_set_111) when allowing for 0 base pair mismatches (i.e. exact matches only). Allowing for 1 base pair mismatch yielded a single hit to an IS3 family IS*Pg*5 transposase ORF A in two ICEs (GCA_018141745.1_ICE8/GCA_018141765.1_ICE9) encoded by two derivative sub-strains of the same *Pg*, suggesting the transposase itself is targeted rather than the ICE. In other bacterial species ICEs have been identified as subjects of CRISPR defenses^44–47^, and thus the ubiquitous nature of ICEs in *Pg*, coupled with the lack of CRISPR defenses against them, suggests that the ICEs may either efficiently attenuate the ability of *Pg* to protect against them or that they are preferentially maintained.

To better understand the nature of the two dominant groups of ICEs in *Pg*, and how they might be differentially shaping *Pg* gene repertoires, we next focused our investigation on genes in Group 1 and Group 2 ICEs. To define the set of high confidence full length ICEs for studies of gene content we used several orthogonal curation approaches developed in our previous work with the *Pg* prophages^14^. We consider the mobile element-specific predictions (here, the ICEfinder^31^ predictions) in light of bacterial species pangenome annotations (which highlight mid- and low-frequency genes in *Pg*, often associated to mobile elements including phages and ICEs), all-by-all *Pg* genome BLAST-based genomic comparisons (often revealing sharp boundaries of inserted elements), and inspecting candidate termini to identify bounding repeat motifs (Fig. 3). This approach yielded 89 fully curated ICEs (*Pg*_ICE_pilot_set_89, curated from 143 total ICEs identified in *Pg*_set_79, Supplementary Data File 1).

**Figure 3.**
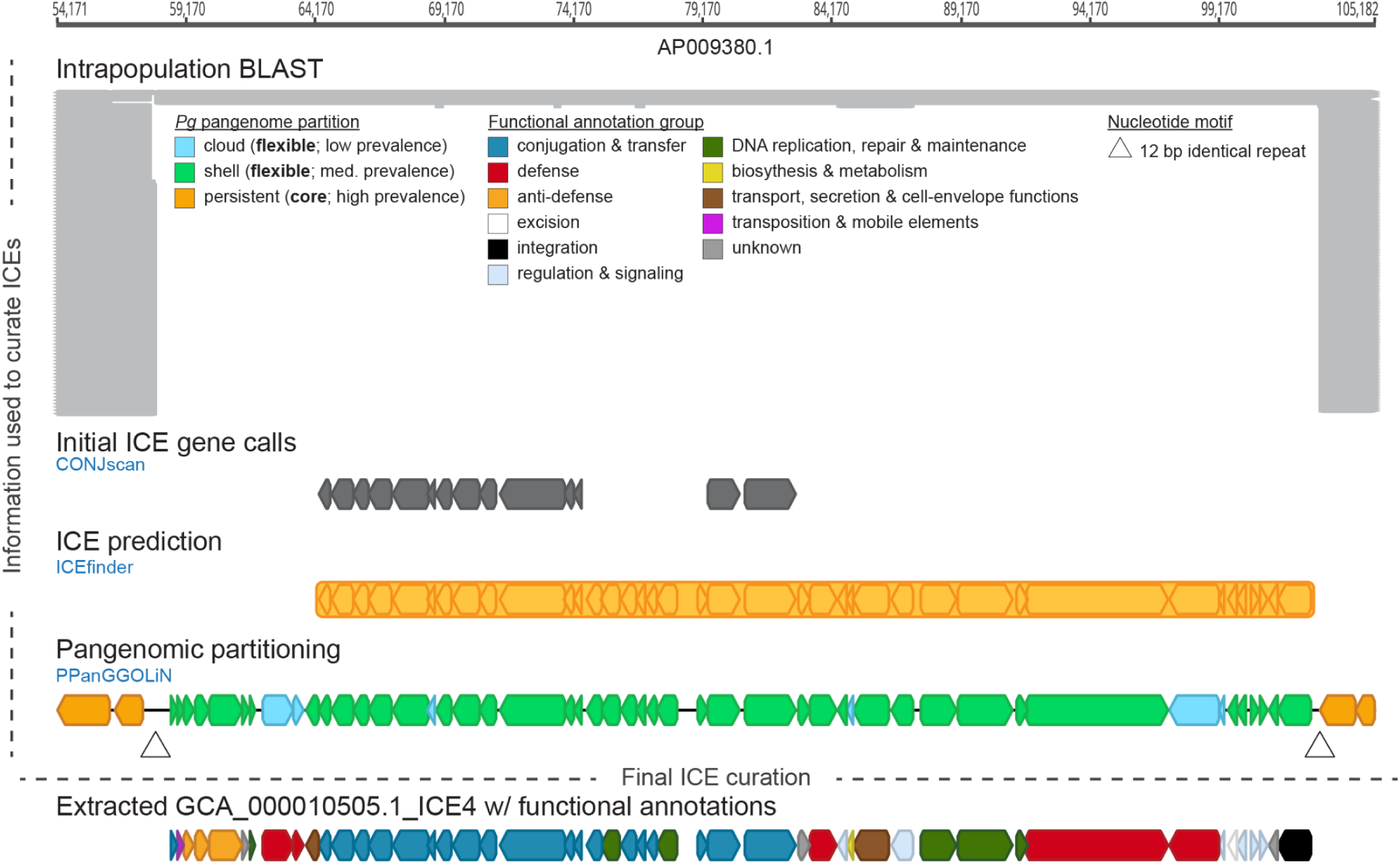
Complementary bioinformatic approaches enabled curation of complete ICEs from the *Pg* genomes. An adapted example view of a region of *Pg* contig AP009380.1 (GCA_000010505.1; ATCC 33277) from Geneious bioinformatic software highlighting the various lines of evidence used in the manual ICE curations. Genes predicted to be part of a conjugative system were initially identified with CONJscan^30^ (indicated by dark gray block arrows). Putative ICE regions were predicted with ICEfinder^31^ (indicated by yellow bar). Pangenomic partitioning was performed by PPanGGOLiN^48^ which identifies whether genes are part of *Pg*’s flexible (light green and cyan block arrows) or core (orange block arrows) pangenome. Repeated nucleotide sequences were identified by the Geneious Repeat Finder (indicated by triangle). All-by-all intrapopulation BLAST searches used to compare each *Pg* genome against all other *Pg* genomes revealed regions that lacked conservation (indicated by gray bars). The final curated ICE (GCA_000010505.1_ICE4) was manually extracted with the aid of all of these analyses.

We combined protein function annotation with pangenome analyses of each of the ICE groups, to learn about conserved and variable genes within each of the groups. Open reading frame predictions were inherited from Bakta^49^ annotations of parent scaffolds and we complemented the Bakta^49^ annotations with additional sequence-, structure-, and protein language model-based annotations (see “Materials and Methods”). To maximize the number of genes to which we can assign function, we clustered all proteins into families (≥90% query and subject cover and ≥70% percent identity) and manually reviewed annotations for all families, assigning a single harmonized annotation (and broad category) to all members of each gene family (see Supplementary Data File 1 for all annotations). We then carried out a pangenome analysis for each ICE group to determine which gene families are conserved within each group and which are accessory (i.e. medium-frequency, and low-frequency).

Overall, we found that both ICE groups exhibit a backbone of conserved hallmark conjugation and mobilization genes (Fig. 4A,B, orange nodes represent gene families present in the majority of ICEs in each group), as well as distinct populations of accessory genes (Fig. 4A,B light green medium-frequency genes, cyan low-frequency genes) often occurring in specific “islands” representing distinct functions. Both groups of ICEs encode notable anti-defenses, defense systems, and potentially beneficial cargo genes, which are highlighted and discussed in detail below.

**Figure 4.**
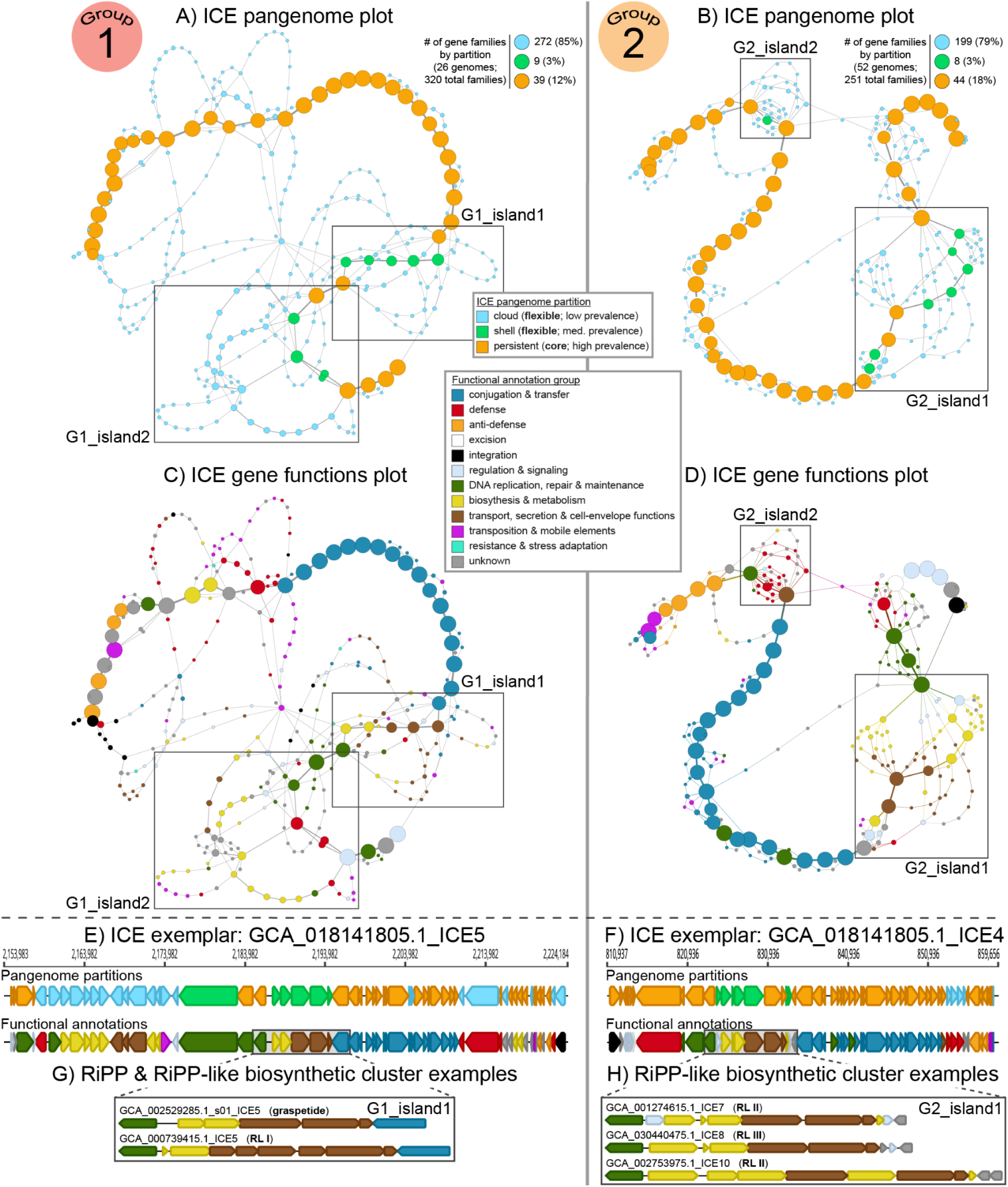
Group 1 and Group 2 *Pg* ICEs show characteristic conserved genes and functional islands. Genome graph representations of each ICE group highlight element backbone structure and gene neighborhoods, with each node representing a protein-coding gene family (≥90% query and subject cover, ≥70% amino acid identity), node size representing the number of genomes in which it is found in the ICE group, and edges representing gene-adjacency; pangenome graphs for Group 1 and 2 represent 26 and 52 curated ICEs, respectively (A, B, C, D). Coloring nodes with group-specific ICE pangenome partition information (i.e. gene partitioning was determined in regards to ICEs within the same group) highlights genes that are conserved within each group of ICEs (orange), present at intermediate frequency within each ICE group (light green), or present in few ICE genomes in that group (cyan) (A, B). Coloring nodes of the genome graphs with predicted protein functions highlights conserved and variable functions within the groups (C, D). Genome diagrams for an exemplar ICE from each group highlight the correspondence between the group-level graphs and individual genomes; note that both exemplars are from strain GMU202011 (GCA_018141805.1), reported as isolated from pus of a patient with severe pneumonia (https://www.ncbi.nlm.nih.gov/nuccore/2030184694) (E, F). Vignettes highlight examples of distinct sets of RiPP and RiPP-like (RL) biosynthesis and transport system genes encoded by different ICEs in each group (G, H). Group 1 ICE vignettes include a graspetide biosynthetic gene cluster (top gene diagram in G; note that precursor is not called by the standard gene caller) and a DKAHGB_00252 gene family precursor RiPP-like system (bottom gene diagram in G, “RL I” in Fig. 5). Group 2 ICE vignettes include a AAIIIG_00105 precursor RiPP-like system (top and bottom gene diagram in H, “RL II” in Fig. 5) and RiPP-like systems with lanthipeptide B-like and novel precursors (middle gene diagram in H, “RL III” in Fig. 5). All genes represented in the figure are highlighted in Supplementary Data File 1.

### Pg ICEs commonly encode anti-defense genes, which are conserved within groups

Recent studies have shown that conjugative elements are enriched in anti-defense systems that promote their survival on entry into new recipient cells^50^, and we find that anti-defense systems are commonly present as conserved genes in *Pg* ICEs. Specifically, we find that 85% of ICEs in Group 1, and 100% of ICEs in Group 2, encode anti-CRISPR protein genes. In addition, both groups encode proteins with predicted similarity to characterized DNA-mimic proteins, although these belong to distinct classes (DMP19 in Group 1, ArdA in Group 2); of these, the ArdA family has an anti-defense function.

The AcrIIA9 anti-CRISPR protein (of which homologs are found in both Group 1 and Group 2 ICEs) has been shown to inhibit Cas9 nuclease activity of the Type II-A CRISPR-Cas system, preventing effective binding and cleavage of targeted DNA^51^. Originally identified within metagenomic libraries from the human gut, AcrIIA9 homologs are found in over 300 species, predominantly within the phylum *Bacteroidetes*^51^. Although these proteins were originally proposed to be of viral origin because many were found in predicted phage sequences^51^, our findings suggest that AcrIIA9-like proteins also contribute to ICE ecology. As our previous work^14^ characterizing the CRISPR-Cas systems in *Pg* did not reveal the presence of type II-A systems, it remains unclear to what extent these AcrIIA9 systems would inhibit the CRISPR systems in Pg.

DNA-mimic proteins, such as DMP19 (of which a potential remote homolog is found in 100% of Group 1 ICEs, as inferred from annotation of DUF4375^52^) and ArdA (found in 94% of Group 2 ICEs), have diverse functions. Whereas DMP19 has thus far been shown to play a role in gene regulation^53,54^, ArdA has been shown to play a role in both regulation and anti-defense, binding to Type I restriction-modification (RM) enzymes and preventing them from binding to and cleaving target DNA^55^. ArdA proteins provide attenuation of RM cleavage within a variety of MGEs, including conjugative plasmids^56^, phages^57^, and ICEs^55^. Our previous analyses characterizing the diversity of defense systems in *Pg* found that Type I RM systems are prevalent^14^, suggesting that these ICE-encoded ArdA proteins may function as DNA mimic decoys in *Pg*, ultimately allowing for ICE transfer. However, although prevalent, Type I RM systems are not ubiquitous across *Pg* strains^14^ and some ICEs found in *Pg* strains that lack a Type I RM system encode an ArdA protein (e.g. ATCC 33277).

### Pg ICEs commonly encode defense system genes, which vary within groups and between elements

Mobile genetic elements have been recognized as “guns for hire”^15^, enriched in defense systems acting against phages and other mobile elements, and we find this to be true also of *Pg* ICEs. Notably, in contrast to anti-defense genes, which are represented by only a limited number of conserved gene families in each ICE group, defense systems are diverse, comprising numerous gene families, each found in only a small proportion of ICEs (Fig. 4A-D, light blue cloud gene families, with red defense functions).

We previously established that *Pg* strains encode diverse defense systems^14^. To evaluate the contribution of ICEs to these defense gene repertoires we used DefenseFinder to annotate ICEs and found that nearly half encode at least one defense system with predicted anti-phage activity (49%; 44/89 in *Pg*_ICE_pilot_set_89), while 18% (16/89) of ICEs encode multiple (Fig. 5). Toxin-antitoxin (TA) systems were the most abundant class of defenses, found in 27% of ICEs overall (24/89), with the SanaTA^60^ system predominant in Group 1 (73%; 19/26) and the ShosTA^61^ in Group 5 (67%; 2/3). TA system “addiction modules” include a stable toxin and an unstable cognate antitoxin, requiring the gene encoding the antitoxin (here, in the ICE) to be maintained in the host cell to ensure activity of the antitoxin and prevention of cell death. TAs have been shown to play an essential role in maintaining ICE stability during vertical passage^62^, yet also to contribute to defense during phage infection by a variety of mechanisms^63–65^. A second class of defense systems common in the *Pg* ICEs is restriction-modification (RM)^66^, with RM systems found in 15% (13/89). The ICE groups differ in the RM system types they predominantly encode, with Group 1 ICEs having type IIG RM systems (23%; 6/26), Group 2 ICEs having type II^67^ (8%; 4/52), and Group 5 ICEs having type IIG (67%; 2/3) and III (33%; 1/3). RM systems function in defense by cleaving phage DNA and effectively shutting down phage replication and integration. A third class of common defense genes identified was that of abortive infection (Abi) systems, found in 15% (13/89) of ICEs. Again, the ICE groups differed in the types of Abi they encoded, with AbiD^68^ found in Group 1 ICEs (27%; 7/26), Group 3 ICEs (50%; 1/2), and Group 4 ICEs (100%; 1/1), AbiE^68^ and AbiJ^69^ found in Group 5 ICEs (100%; 3/3) and (33%; 1/3), respectively. No Abi systems were identified in Group 2 ICEs. Abi systems act by killing the host cell upon sensing of phage infection, thereby providing population-level immunity by protecting neighboring cells. Other lower frequency systems, all with predicted anti-phage functions, including Thoeris II^70^ in Group 2 (13%; 7/52), NLR-like bNACHT0960 in Group 1 (8%; 2/26), SspBCDE^71^ in Group 1 (8%; 2/26), PD-T7-2^72^ in Group 1 (4%; 1/26), and VP1853 in Group 2 (2%; 1/52), were identified.

**Figure 5.**
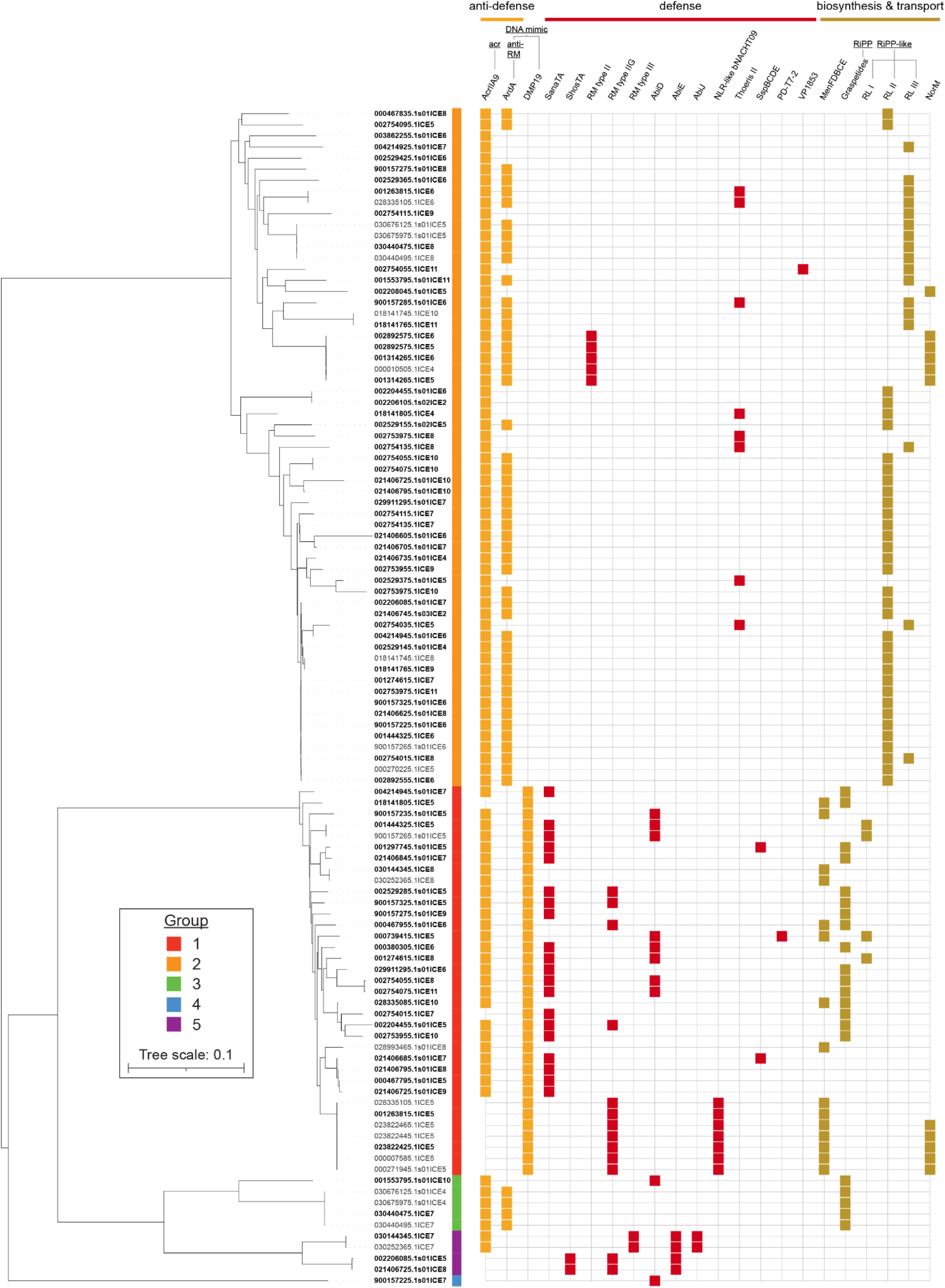
*Pg* ICEs encode proteins with functions that shape the anti-defense, defense, and metabolic repertoires of their hosts. Proteomic tree (midpoint-rooted) of *Pg* ICEs generated with ViPTree^58^, based on proteome similarity of ICE conjugation modules (*mobC*-*traQ*; grouped and colored by 70% nucleotide similarity of conjugation modules determined by VIRIDIC^33^). All ICEs from *Pg*_ICE_pilot_set_111 for which conjugation modules could readily be identified and extracted were included (106 total), ICEs from the non-redundant *Pg*_ICE_pilot_set_89 are bolded. Presence of anti-defense, defense, and biosynthesis and transport systems in each ICE are denoted by orange, red, and mustard-colored boxes, respectively. The mechanisms of each anti-defense system are denoted: anti-CRISPR (acr) for AcrIIA9 and anti-restriction (anti-RM) for ArdA; ArdA and DMP19 are both indicated as DNA mimic proteins, while only ArdA has a confirmed anti-defense function. Defense genes include those defined by DefenseFinder^59^ and thus represent a conservative survey given the challenges around annotating such systems, for example, we note that the diverse putative novel defense systems present in the “defense island” in Group 2 ICE genomes are not represented here. RiPP and RiPP-like (RL) systems are presented as four major groups based on distribution, including graspetide biosynthetic gene clusters (in G1 ICEs), DKAHGB_00252 gene family precursor peptide systems (RL I; G1 ICEs), AAIIIG_00105/AIIIG_00110 systems (RL II; G2 ICEs), and a pool of various other, lower frequency groups (RL III) found in G2 ICEs and representing novel and lanthipeptide B-like precursors (BAOJBJ_02091, ABJECD_01646, GDPKEO_01401, GHBHDG_01816, JICNJJ_00670, KBLCOE_01182, MCAPDE_01026). An interactive version of the iTOL^34^ tree can be found online at https://itol.embl.de/tree/70177211213255761785532678.

Overall, we find that ICE groups show distinct patterns of common defenses they encode, yet individual ICEs are unique in the specific defense gene families they encode and thereby confer on their bacterial host strains. Of note, in the aforementioned analyses, we exclusively consider defense systems annotated by DefenseFinder, however, novel systems continue to be identified and described and many genes of unknown function may ultimately function in defense, thus our representation of defenses in ICEs is conservative.

Notably, for example, Group 2 ICEs show a distinctive “defense island” region of their genome (Fig. 4B,D, G2_island_2), into which genes conferring defense functions are commonly found inserted - genes of unknown function in these islands likely represent novel defense systems waiting to be investigated.

### Pg ICEs commonly encode biosynthetic gene clusters, which differ between groups

Evaluating gene functions in the *Pg* ICEs revealed that these mobile elements commonly encode genes associated with putative biosynthetic gene clusters (BGCs). Particularly notable are the menaquinone synthesis pathway genes (Fig. 6) and diverse ribosomally synthesized and post-translationally modified peptides (RiPPs) system candidates and their associated putative transport systems (“graspetide” and “RiPP-like system candidates” in Fig. 5). We highlight these further below.

**Figure 6.**
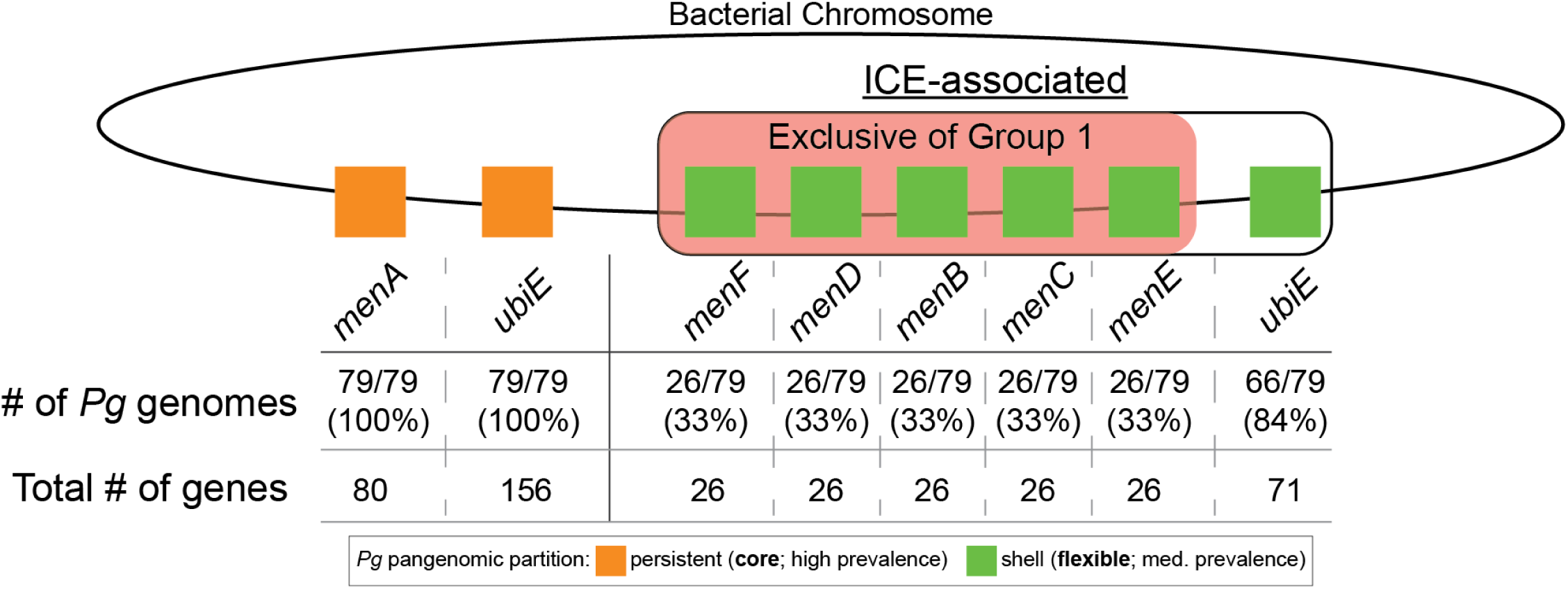
Menaquinone pathway genes *menFDBCE* are associated with ICEs in *Pg*. Schematic diagram exemplifying the differences in association of genes involved in *Pg*’s menaquinone synthesis pathway with the bacterial chromosomal backbone or ICEs. Total number of *Pg*_set_79 genomes each gene is present in, at least once, is denoted, as well as the total number of each gene within those genomes. Genes are shown as squares (colored based on their pangenomic partition) located in either the bacterial chromosomal backbone or ICE region based on their presence (or lack thereof) in curated ICEs, near ICE conjugation module regions, or in ICE-like regions (see “Materials and Methods”).

Menaquinone, an essential electron carrier in anaerobic respiration, is a vital growth factor for oral anaerobes who cannot synthesize it themselves. Many strains of *Pg* lack the full pathway for menaquinone synthesis, for example, relying on exogenous addition of 1,4-dihydroxy-2-naphthoic acid (DHNA), menaquinone, menadione, or phylloquinone, for growth in laboratory cultures^73–75^. A recent study^73^ of growth factors enabling cultivation of fastidious oral bacteria observed that a 5-gene *menFDBCE* operon is either entirely present or absent in oral *Porphyromonas* and *Tannerella* species, whereas two genes in the pathway, *menA* and *ubiE*, are universally encoded as individual genes elsewhere in the genomes. Additionally, they noted that 31% of *Pg* genomes investigated contain 7 genes of the pathway (excluding *menH*), and that the presence of this operon was associated with ability to grow without addition of menadione or DHNA to the media. Here, we similarly find that 33% (26/79) of *Pg* strains encode the full pathway, but further observe that in all cases where *Pg* strains do encode the variable *menFDBCE* operon, it is always present on a Group 1 ICE (Fig. 6). Therefore, it is possible that the ICEs found in *Pg*, and other species, encoding this locus could confer a benefit to their host by enabling them to produce endogenous menaquinone, reducing their reliance on nearby growth factor-producing neighbors.

With regard to the observed RiPP and RiPP-like systems, we observe that each group of ICEs has its own distinct genomic island “hot spot” harboring various putative RiPPs, and that the two groups differ markedly in the classes they encode (Fig. 4G,H, Fig.5). Group 1 ICEs are distinct in their carriage of graspetide BGCs, a class of RiPPs noted as being present in *Pg*^76,77^, however not recognized as encoded on mobile elements. Graspetides are a class of RiPP in which a precursor peptide undergoes ATP-grasp-mediated macrocyclization via ester or amide side chain linkages^78^. All functionally characterized graspetides have been shown to be protease inhibitors, with the serine protease-inhibiting microviridins among the most well-characterized examples. In the *Pg* ICEs, the graspetide BGCs are defined by the presence of two grasp-with-SPASM biosynthesis genes (*gwsG*/*gwsS*) and these are enriched in Group 1 (58%; 15/26) and Group 3 (100%; 2/2) ICEs, respectively, and are present in 21% (19/89) of ICEs overall in *Pg*_ICE_pilot_set_89 (Fig. 5). Notably, the essential precursor protein is often not called and thus requires special curation to be detected; we find that the ICE-encoded grasp-with-SPASM operon features a 222 bp propeptide immediately upstream of the GwsG and GwsS ligases. In addition, these loci encode a secretion apparatus resembling a canonical type I secretion system (outer membrane β-barrel protein, ABC transporter, and HlyD-like membrane transporter) immediately downstream, consistent with secretion of the synthesized graspetide.

Group 2 ICEs do not encode graspetide BGCs, however also have a region of their genomes where RiPP-like genes are commonly found (Fig. 4H “G2_island_1”). The majority of Group 2 ICEs encode genes representing a potential novel thiopeptide-like RiPP biosynthetic gene cluster, with others harboring alternative sets of biosynthetic and transport genes together with short proteins identified as candidate RiPP precursors, including lanthipeptide B-like examples, per predictions with RiPPMiner^79,80^.

We also found that ICEs in Group 1 (4%; 1/26 Pg_ICE_pilot_set_89) and Group 2 (10%; 5/52) encode the multidrug efflux pump NorM (Fig. 5). The NorM efflux pump subfamily belongs to the MATE (multidrug and toxic compound extrusion) family and is well studied in Gram-negative bacteria such as *Vibrio* species and *Neisseria gonorrhoeae* where it is recognized for its ability to export antibiotics (e.g. fluoroquinolones), cationic dyes, and other toxic compounds^81,82^. Finding this efflux pump encoded in *Pg* ICEs raises the possibility that *Pg* harboring these ICEs may be positively selected for as a result of antibiotic dental treatment, contributing to the increase in antibiotic resistance in *Pg*^83^. Similar to the other systems discussed, these findings suggest that genes encoded by ICEs may provide ICE-containing *Pg* strains a competitive advantage in the stressful environment of the subgingival crevice.

### Pg ICE groups differ in their distributions across taxa in the oral microbiome

The observations of the prevalence and functional potential of *Pg* ICEs are striking and raise the question of their potential to spread traits across taxa beyond *Pg*. As mentioned above, recent work demonstrated that an ICE region in ATCC 33277 (the canonical CTnPg1, and here defined as a Group 2 ICE) was detected in oral metagenomic samples even in the absence of *Pg*, and was associated with diverse host species. To understand these findings within the framework of our newly classified ICE groups, we investigated the distribution of the representative TraG gene as a proxy for each ICE group in a search of all oral genomes in the *e*HOMD^29,84^. We find that Group 1 ICE TraG sequences are found predominantly in *Pg*, as well as in fellow ‘red complex’ periodontal pathogen *Tannerella forsythia,* and *Bacteroides heparinolyticus;* including, for example, a shared high identity ICE region present in both *Pg* MP4-504 (contig LOEL01000078.1) and *Tf* ATCC 43037 (JUET01000075.1). By contrast, and consistent with Torres-Morales’ findings^26^, Group 2 ICE proxy TraG genes show wide distributions across *Porphyromonadaceae*, *Prevotellaceae*, as well as are found in species in the *Atopobiaceae*, *Bacteroidaceae*, and *Tannerellaceae* (Fig. 7).

**Figure 7.**
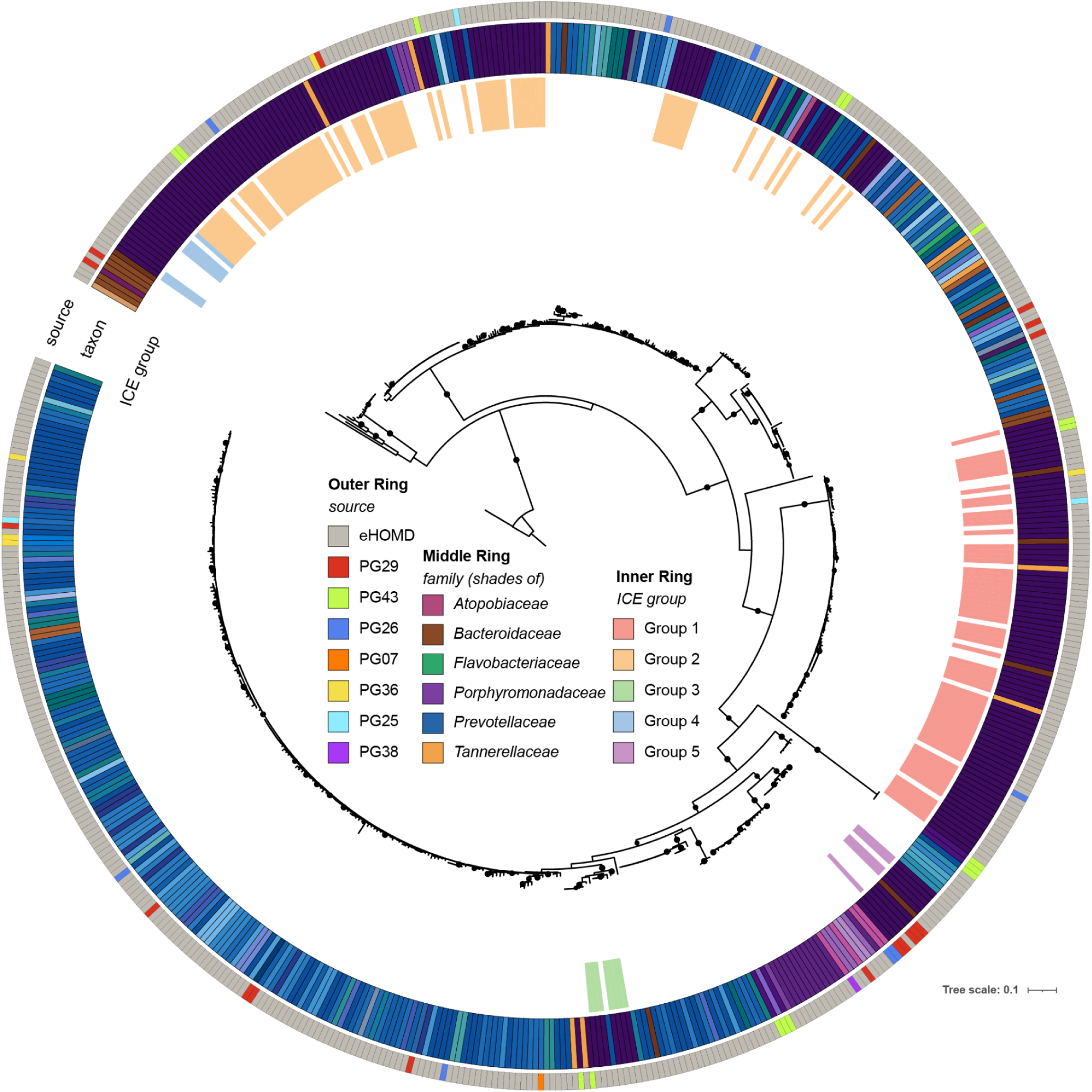
*Pg* ICE-like TraG genes show distinct taxonomic distributions across bacterial taxa in the *e*HOMD and clinical samples. To infer the presence of related ICEs in *e*HOMD and clinical samples, a characteristic ICE marker gene protein sequence, TraG, was used as a query in BLASTP-based searches; the *Pg* W83 Group 1 ICE TraG (eHOMD^29^ Prokka^87^ ID: GCA_000007585.1_01362) was the representative query and all shown sequences are ≥99% query coverage ≥65% sequence identity. Outermost ring indicates sequence source: searches were performed against all bacterial genomes in the *e*HOMD (grey indicates *e*HOMD oral taxon) and against contigs assembled from enrichment culture mini-metagenomes from clinical donors with periodontal disease (neon colors indicate donor ID). Middle ring indicates predicted contig taxon: leaves representing *e*HOMD genomes are colored by known source taxon, leaves representing metagenomic sequences are colored by taxon predicted by Kraken 2^86^ for the contig on which the TraG is encoded. Innermost ring indicates group classification for leaves representing sequences encoded within *e*HOMD *Pg* ICE conjugation modules. Bootstrap support of SH-aLRT ≥80% and UFboot ≥95% indicated with filled black circles. The lower-identity protein sequence of *Phocaeicola vulgatus* (GCA_024759565.1_00679) was used as the outgroup. An interactive version of the iTOL^34^ tree can be found online at https://itol.embl.de/tree/70177211213457111785147405 and shows full protein IDs and the ability to follow hyperlinks to JBrowse views of genomic neighborhoods on the *e*HOMD webserver. Underlying data available in Supplementary Data File 2.

#### Individuals may harbor multiple distinct Pg ICEs in their oral microbiome

To understand the extent to which ICEs contribute to strain-level diversity in *Pg* within individuals, we investigated subgingival plaque samples from our recent longitudinal clinical study of donors with periodontal disease and qPCR-confirmed *Pg*-carriage (see “Materials and Methods”). To overcome the challenge of detecting MGEs in *Pg*, which is expected to occur at low relative abundance and thus pose challenges to assembly from direct metagenomic sequencing approaches, we developed a novel strategy combining enrichment-based cultivation of *Pg* with a deep-sampling enrichment-cultivation mini-metagenome approach (*Pg*-ECMM, see “Materials and Methods”). Using this approach, we investigated the presence and distributions of *Pg* ICEs in our clinical samples. We generated *Pg*-ECMM datasets for subgingival samples from 8 study subjects, including pooled quadrant 1 plaque for all 8 donors, as well as pooled quadrant 3 plaque from an approximately 6-month later timepoint for a subset of 4 donors. To identify ICEs, we assembled each of the *Pg*-ECMMs for each donor using MEGAHIT^85^, annotated the contigs as to which species they were derived from using Kraken 2^86^, and searched all contigs for TraG genes using the same query as used for the *e*HOMD searches above.

We found that representatives of four ICE groups were detectable via *TraG* marker genes in subgingival plaque of recent clinical samples collected from our *Pg*-positive donors (Fig. 7, leaves on the tree marked with a neon-colored color strip in the outermost ring are from donor samples). Altogether, this approach detected *Pg* ICEs in 6 of 8 subjects, with 4 individuals predicted to harbor at least two distinct ICE groups based on marker gene fingerprints, commonly Group 1 and Group 2, consistent with our observations from studies of historical isolate genomes. One of the subjects sampled longitudinally harbored 5 distinct *Pg* ICE TraG sequences, with 4 of these sequences detected at both time points tested (Supplementary Data File 2).

## DISCUSSION

This work identifies ICEs as a pervasive feature of *Pg* genome ecology and physiology, with substantial potential to shape the success and interactions of this pathobiont in the oral microbiome. We build on the foundational work of Naito et al.^18,19^ and Tribble et al.^27^, who first identified and characterized ICEs in *Pg* with their studies of the *Pg* ATCC 33277 CTnPg1, which we here classify as a Group 2 ICE. Our findings shed light on a recent study by Torres-Morales et al.^26^, who showed, by mapping reads from oral metagenomes, that the *Pg* ATCC 33277 Group 2 ICE is present in oral microbiomes even in the absence of *Pg*. They found that genomic backgrounds recruiting reads from the ATCC 33277 ICE belonged to diverse other species in the Bacteroidales, and this was consistent with our findings of the broader distributions of Group 2 ICEs across these same species in our clinical enrichment culture collection. Together, these previous studies and our new findings offer a new model for understanding aspects of gene flow in *Pg* as being shaped by the ecologies of specific ICEs with distinct gene cargos (as summarized in Fig. 8).

**Figure 8.**
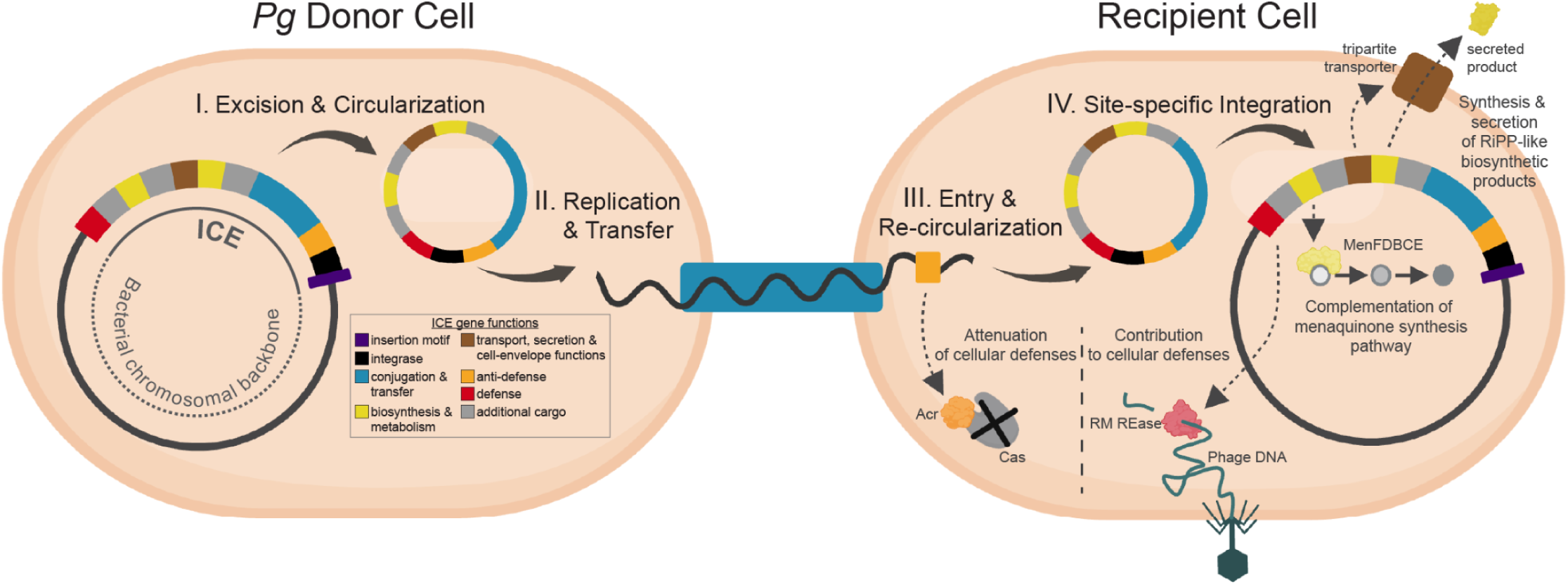
Schematic representation of ICE transfer between cells, with example annotated functions encoded by *Pg* ICEs. Diagram depicts the four main steps of ICE transfer from a donor cell to a recipient cell: ICE excision from the donor’s chromosome and circularization (I), replication and transfer through the ICE-encoded T4SS apparatus (II), entry into the recipient cell and re-circularization (III), and integration into the recipient’s chromosome. Coloration of the ICE genome represents different functions encoded by the ICE. Examples of functions encoded by ICEs include protection against the host bacteria’s defenses upon entry (anti-defenses), protections against phages and other MGEs afforded to the bacterial host once integrated (defenses), and biosynthesis- and transport-related pathways.

### Contribution of cargo genes to the functional capabilities of Pg

This work has provided initial bioinformatic predictions of the potential of ICEs to shape the physiology of *Pg*. We have shown that the menaquinone synthesis pathways, that enable *Pg* to generate key metabolites beneficial to its survival, are associated with Group 1 ICEs. We therefore predict that conjugation of Group 1 ICEs into strains that do not carry them will result in acquisition of menaquinone synthesis capabilities by those recipient strains. In addition, we have identified RiPP and RiPP-like biosynthetic gene clusters, not previously investigated in the oral microbiome, as common cargo on ICEs, with graspetide systems distinctive of Group 1 ICEs. These observations suggest that biosynthetic functional potential may track ICE distributions and host ranges, and raise the question of to what extent biosynthetic functions are associated to mobile elements such as ICEs more broadly in the human oral microbiome.

### Host range determinants of ICEs

The discovery that gene functions of potential interest are tied to mobile elements underscores the importance of identifying molecular determinants of their host ranges. This knowledge will enable prediction of distributions of genes of interest, as well as their potential to spread between strains and species in the oral microbiome. Of note, nearly all ICEs identified in *Pg* harbor predicted anti-CRISPR genes and most Group 2 ICEs also encode a predicted anti-restriction protein (ArdA). These observations are consistent with recent work showing that ICEs frequently encode anti-defense systems that protect them during entry into new recipient cells^50^. Additional features of ICEs that may play roles as determinants of host range include ICE genes encoding transfer apparatus proteins physically involved in binding to cell surface receptors on the outer membranes of potential recipient cells, the availability of preferred insertion site sequences in genomes across taxa, and the compatibility of ICE replication genes with host cell machinery. Whether *Pg* ICEs with host ranges that span multiple species are targeted by CRISPR in these taxa also remains to be evaluated.

### Competition with phages

This study revealed that *Pg* ICEs encode anti-phage defense systems that may prevent phage infection of their host cells. Comparable to the ICEs also encoding anti-defense proteins to combat *Pg* defenses, *Pg* phages likely encode anti-defense systems to combat the defenses encoded on ICEs. This ICE-phage interaction is exemplified in the same *B. subtilis* system that showed an ICE-encoded protein inhibits phage infection^24^. A recent continuation of this study found that co-infecting phages encode a protein that attenuates the abortive infection protein produced by the ICE^88^. Competition between chromosomally-integrated ICEs and prophages can occur after infection as well when vying for insertion sites. ICEs and prophages are then activated, or induced, from the host chromosome, most canonically in response to DNA damage, in order to replicate and spread throughout bacterial populations. Prophage induction leads to lysis of the host cell, which is problematic for ICEs as the integrity of the cell is vital for effective cell-to-cell contact and conjugation. In the *B. subtilis* system, the ICE in a double lysogen, containing both an ICE and prophage, encodes a system that modulates the DNA-damage SOS response by reducing RecA filaments, thus protecting its own ssDNA during replication and transfer, but also limiting prophage induction and phage-mediated cell death^89^. Overall, a more complete understanding of the co-evolving dynamics between ICEs and prophages may shed light on the difficulties in isolating phages for certain species, including *Pg*, and will facilitate development of robust and efficient phage therapeutics.

## CONCLUSIONS

This work establishes a new conceptual framework identifying ICEs as a pervasive force structuring strain-level gene repertoires and horizontal gene transfer in *Pg*. The finding that two groups of ICEs with distinct gene repertoires and host associations are highly prevalent in *Pg* offers a new perspective on the mechanisms underpinning observed distributions of gene functions in periodontal microbiomes in health and disease, as well as on the challenges that have been faced with isolation of phages for this species. Future studies of the factors shaping the structure and outcomes of these multipartite interactions between *Pg* and its diverse infecting ICEs and phages are predicted to be fruitful in shedding new light on population structure and function in the periodontal niche.

## DECLARATIONS

### Ethics approval and consent to participate

The clinical study referenced here was approved by the Institutional Review Board at the University at Buffalo (IRB number STUDY00005287).

### Availability of data and material

Additional data associated with this study is available through Zenodo (https://zenodo.org/records/21723056), including: 01.Pg_Bakta_gbffs [GenBank format files for Bakta-annotated Pg genomes (for original complete assemblies and scaffolds generated in this work)]; 02.PgICE_cores_fastas [nucleotide FASTA format files for the 160 extracted ICE conjugation module regions]; 03.curated_PgICE_fastas [nucleotide FASTA format files for the curated and extracted ICEs in *Pg*_ICE_pilot_set_111]; 04.curated_PgICE_gffs [gff format files for the curated and extracted ICEs in *Pg*_ICE_pilot_set_111, including Bakta annotations]; 05.curated_PgICE_gbs [GenBank format files for the curated and extracted ICEs in *Pg*_ICE_pilot_set_111, including Bakta annotations]; 06.PgICE_G1_G2_pangenomes [PPanGGOLiN analyses output files for Group 1 and Group 2 ICEs in *Pg*_ICE_pilot_set_89].

## Authors’ contributions

CBM - conceived and designed the study, developed and executed methods for identifying and characterizing

ICEs, contributed to laboratory isolation of strains, wrote the original draft of the manuscript

EMH - developed and optimized methods for extraction and sequencing of mini-metagenomes, read and gave feedback on the manuscript

AKM - contributed to annotation of ICEs, read and gave feedback on the manuscript SS - conducted donor screening and carried out clinical study

DS - assisted with clinical study and sample collection, read and gave feedback on the manuscript MS - assisted with screening of donor samples, read and gave feedback on the manuscript

PID - designed the clinical study, read and gave feedback on the manuscript

KMK - contributed to conception and design of the study, bioinformatic analyses, laboratory studies, and writing of the manuscript

## Acknowledgements

We thank the donors to our clinical study for making these findings possible. We thank Takuma Suzuki for guidance in sample processing, as well as other members of the Diaz lab who assisted with sample collection and screening. We thank Ashu Sharma, Michelle Visser, Chelsie Armbruster, and Libusha Kelly for helpful discussions and feedback on the work. Compute resources enabling this work were provided by the Center for Computational Research at the University at Buffalo.

## Supplementary Information

**Additional file 1: Supplementary Data File 01. Overview of ICEs and associated protein annotations.** Sheet 01.ICE_summary_table provides an overview of information on each of the 170 ICE regions referenced in the manuscript, including information on the *Pg* strain of origin, its predicted VIRIDIC genus, species, its inclusion in various described sets, and length and %GC. Sheet 02.ICE_cores_map2homd and 03.ICE_curateds_map2homd provide start and stop coordinates for each ICE conjugation module region (170) and curated ICE (111) with respect to contigs available on *e*HOMD, including with hyperlinks to enable ready viewing of these regions with JBrowse. Sheet 04.MEGA_annos provides a synthesis of protein-level annotations for all proteins in the set of 111 curated Pg ICEs, including assignment to gene families as described in “Materials and Methods”. Additional sheets provide more complete annotation outputs from various tools employed for annotation, as described in “Materials and Methods”.

**Additional file 2: Supplementary Data File 02. Data underlying TraG analyses presented in** **Fig. 7**. Sheet 01.BLAST.TraG_vs_clinical provides information on TraG sequences identified in *Pg* enrichment culture mini-metagenomes (*Pg*-ECMMs) from clinical samples. Shown are BLAST results and additional annotations indicating the predicted taxon for the contig on which each sequence was found. Sheet 02.clinical_source_data provides information on source samples used to generate each *Pg*-ECMM. Sheet 03.BLAST.TraG_vs_eHOMD provides information on eHOMD TraG sequences shown in the tree. Shown are filtered BLAST results and additional annotations about source species and taxon Body Site associations. Sheet 04.eHOMD_JBrowse_links provides links to JBrowse views of each *e*HOMD TraG sequence shown in Fig. 7.

## Materials and Methods

### *Pg* genome scaffolding, annotation, and selection

All *Pg* sequences available in *e*HOMD (https://homd.org) were downloaded on January 12^th^, 2026 (Genomic RefSeq v11.02) to be used in these analyses. Of these 99 *Pg* genomes, 36 were complete assemblies and 63 were incomplete. In order to improve fragmented genome assemblies which could hinder the discovery of ICEs, each incomplete genome was scaffolded against each of the 99 *Pg* genome assemblies, in an all-by-all manner, with reference-based scaffolding tool Multi-CSAR^90^[default parameters] (https://github.com/ablab-nthu/Multi-CSAR). Gene calls and annotations were performed for each scaffolded and already complete assembly with Bakta^49^ v1.11.4 [default parameters; except skipped CRISPR arrays annotations due to known bug preventing the annotation of some assemblies] (https://github.com/oschwengers/bakta). To aid in the selection of scaffolded assemblies with the most complete ICEs, conjugative and mobilizable elements were identified with the CONJScan^30^ [default parameters; used ordered replicon database type] (https://github.com/macsy-models/CONJScan) model of MacSyFinder^91^ v2.1.4 (https://github.com/gem-pasteur/macsyfinder). Scaffolded assemblies for *Pg* with originally incomplete assemblies were then selected for downstream analyses based on a tiered ranking system: 1) most complete conjugative systems, 2) highest average score of each conjugative system, 3) fewest number of total scaffolds in the assembly, 4) highest percent of pairs recovered in the scaffold assembly compared to original assembly, 5) highest combined number of MOB systems and decayed conjugative systems. Moving forward, two sets of *Pg* genomes were defined for subsequent analyses: *Pg*_set_99 and *Pg*_set_79. *Pg*_set_79 represents a non-redundant subset of complete and scaffolded *Pg* assemblies from *Pg*_set_99 that was generated by removing re-sequenced strains, sub-strains, and metagenomic assemblies.

### ICE and IME prediction

The complete and scaffolded *Pg* assemblies from *Pg*_set_99 yielded a total of 206 contigs. GenBank files of these contigs produced by Bakta^49^ were run individually through the ICEfinder 2.0^31^ [default parameters; with input GenBank marked as single genome] webserver (https://tool2-mml.sjtu.edu.cn/ICEberg3/ICEfinder.php) that utilizes the ICEberg 3.0 database to predict entire ICEs and IMEs. The ICEfinder^31^ results for each contig were scraped from the output webpage using a Bash script that queried the corresponding job ID.

### ICE grouping and phylogeny

A total of 170 ICEs were predicted in the 206 contigs produced from the *Pg*_set_99 scaffolding. To determine the relatedness of these ICEs, the nucleotide sequence of their conjugation modules that contain both MOB and MPF systems, spanning from gene *mobC* to *traQ*, was extracted in the Geneious^92^ bioinformatic software. This approach, which is similar to what was done in other studies^35–37^, was used because conjugation modules are the most conserved feature of ICEs. Nucleotide sequence similarity comparisons were performed on the extracted conjugation modules with the VIRIDIC^33^ [default parameters] webserver (https://rhea.icbm.uni-oldenburg.de/viridic) and placed into “genus-” and “species-level” OTUs by adopting the respective 70% and 95% nucleotide identity thresholds from the International Committee on Taxonomy of Viruses (ICTV) taxonomy framework for phages^38^. ICEs that fell into the same “genus” based on the sequence similarity of their conjugation modules were simply defined as being in the same “group” for this study. This resulted in 160 ICEs being assigned to one of five distinct groups, with 10 not assigned to any group. Conjugation modules from exemplar ICEs and their gene clustering was visualized with Clinker^41^ v0.0.31 [default parameters; except using a minimum 69% amino acid identity threshold which maintains *mobC* and *traG* clustering together between groups] (https://github.com/gamcil/clinker). The phylogenetic relationship of ICEs was resolved by generating a proteome-based similarity tree of the grouped ICEs by submitting their conjugation module nucleotide sequences to the ViPTree^58^ [default parameters; except analysis was run to only include the query] webserver (https://www.genome.jp/viptree) which calls proteins with Prodigal^93^. Negative tree branch lengths were manually set to zero. The resulting midpoint-rooted ICE proteomic phylogenetic tree was visualized with iTOL^34^ (https://itol.embl.de).

### ICE curation

To establish a dataset of high-confidence, complete ICEs, manual curation was performed to refine the predictions by ICEfinder^31^. This complementary approach was similar to what was performed when curating prophages in *Pg* genomes^14^. In combination with the ICE predictions by ICEfinder^31^ and groupings by VIRIDIC^33^, all the following information was visualized in Geneious^92^ for each *Pg* assembly to help make precise nucleotide-level boundary calls of ICE regions: pangenomic partitioning for each gene by PPanGGOLiN^94^ v2.2.6 [default parameters] (https://github.com/labgem/PPanGGOLiN); identification of plasmids, in case an ICE was cross-annotated as a plasmid, by geNomad^95^ v1.11.2 [default parameters] (https://github.com/apcamargo/genomad); all-by-all sequence comparison for each contig by BLAST^96^ v2.17.0 [default parameters; except set 98% nucleotide identity threshold and a word size of 100] (https://www.ncbi.nlm.nih.gov/books/NBK279690); and identification of nucleotide repeats by Repeat Finder v1.0.1 in Geneious^92^. This approach resulted in a pilot dataset of 111 fully-curated ICEs, referred to as *Pg*_ICE_pilot_set_111, from *Pg*_set_99 assemblies. A subset of this pilot dataset, referred to as *Pg*_ICE_pilot_set_89, was derived from the non-redundant *Pg*_set_79 assemblies.

### *Pg* phylogeny

To define the *Pg* core genome and generate a core gene nucleotide alignment, Panaroo^32^ v1.5.2 [default parameters; with strict cleaning mode and core alignment] (https://github.com/gtonkinhill/panaroo) was run using Prokka^87^-generated *Pg* GFF files downloaded from HOMD (excluding metagenomic assemblies that could obscure phylogenetic inferences). *Porphyromonas gulae* COT-052 OH3856 (GCF_00765945.1) was also included in this pipeline as the outgroup genome. The core gene alignment was then used to generate a phylogenetic tree with FastTree^97^ v2.1.11 [default parameters; except a nucleotide alignment was run with the GTR nucleotide evolution model] (https://github.com/morgannprice/fasttree). The resulting *Pg* core gene phylogenetic tree was visualized with iTOL^34^ (https://itol.embl.de).

### CRISPR spacer mapping to ICEs

CRISPR spacers encoded in each *Pg* assembly were identified with CRISPRCasTyper^43^ (CCTyper) v1.8.0 [default parameters; using the metagenomic Prodigal^93^ mode] (https://github.com/Russel88/CRISPRCasTyper). To determine if *Pg* CRISPR spacers target ICEs, the spacer sequences were aligned to the ICE sequences by Bowtie^98^ v1.3.1 [default parameters; except allowing for either 0 or 1 mismatch in the target sequence] (https://github.com/BenLangmead/bowtie).

### Extended ICE annotations

Predicted proteins from the curated *Pg*_ICE_pilot_set_111 (*Pg*_ICE_pilot_set_111.all_protein_genes.faa), which comprised 6,495 total sequences, were annotated using multiple complementary approaches (Supplementary Data File 1), as follows. Signal peptides were predicted with SignalP6.0^99^ (fast model; –organism other; –mode fast; –bsize 96; other HPC job resources: 12 CPU and 32G memory). Prokaryotic subcellular localizations were predicted using DeepLocPro^100^ v1.0 (--group any; HPC job resources: 1 GPU (any type), 8 CPU, 32G memory). Gram-negative bacterial secretion system cargo (T1SS, T2SS, T3SS, T5SS, and T6SS) were predicted with DeepSecE^101^ v0.1.2, an ESM-2 based classifier, using the tool’s default configuration that was tested and verified through five separate rounds of data validation (HPC job resources: 4 CPU, 32G memory, 1 GPU). Protein domains and family memberships were identified with InterProScan^102^ v5.76-107.0 using its default member databases (HPC job resources: 20 CPU, 64G memory). All tools were run on the University at Buffalo’s high performance compute cluster at the Center for Computational Research (CCR). SignalP 6.0, DeepLocPro, and DeepSecE ran on NVIDIA GPUs (any available, with the exception of DeepSecE which we constrained to ‘A100|V100|H100’). Note: Certain bioinformatic tools, including DeepLocPro, DeepSecE, and InterProScan require the use of a fasta file lacking the trailing translational stop (*) symbol. Percent GC for each ICE was also calculated using seqkit v2.7.0 (fx2tab -n -g).

Additional webserver-based annotations included the following. Prediction of defense genes with the Prokaryotic Antiviral Defence LOCator^103^ (PADLOC) v2.0.0 with PADLOC-DB v2.0.0 (https://padloc.otago.ac.nz/padloc/), including option for CRISPRDetect, with analyses performed on Geneious-derived genbank files from which clipped terminal genes were removed. Prediction of defense genes also with DefenseFinder^59^ v2.0.0 with Models v2.0.2 (https://defensefinder.mdmlab.fr/), with analyses performed on protein sequences. Annotation of carbohydrate active enzymes with dbCAN3^104^ (https://pro.unl.edu/dbCAN2/). Broad functional annotation using the Galaxy Europe instance of eggNOG-Mapper^105^ (eggNOG Database v5.0.2), and DeepKOALA^106^ (https://www.genome.jp/tools/deepkoala/), with analyses performed on protein sequences. Individual proteins were evaluated using HHpred^107^ with the MPI Bioinformatics Toolkit^108^ (https://toolkit.tuebingen.mpg.de/tools/hhpred) and RiPPMiner^79,80^ (http://www.nii.ac.in/∼priyesh/lantipepDB/new_predictions/index.php).

To synthesize annotations, and assign each to broad functional categories, we used the following approach. First, we assign each protein to a protein family using clustering with MMseqs2^109^ v18.8cc5c [c 0.90, min-seq-id 0.70, cov-mode 0, cluster-mode 0] requiring bidirectional ≥70% identity and ≥90 coverage to assign proteins to the same cluster. Next, we consolidate all annotations, including ICE group- and species-assignments and PPanGGOLiN^94^ partition assignments for Groups 1 and 2 (see below). We curate all available data for each protein and family by coupling iterative manual review and iterative AI-assisted synthesis, yielding 8 generated “ANNO” summary columns (category, tag, short_name, full_name, localization, confidence, primary_evidence, justification_note).

### ICE pangenome analyses

Pangenome partitioning analyses for Group 1 and 2 ICEs were performed with Pg_ICE_pilot_set_89, which includes curated 26 Group 1 ICEs and 52 Group 2 ICEs from non-redundant *Pg* strains. Pangenome analysis was performed using PPanGGOLiN^94^ v2.3.0 [default parameters] separately for the 26 Group 1 ICEs and the 52 Group 2 ICEs, using genbanks with clipped genes removed (as described above) and pre-defined protein family assignments based on the MMseqs2^109^ clusters representing all curated ICE proteins (described above).

Visualization of pangenome graphs was performed as follows. Pangenome graph outputs produced by PPanGGOLiN for Group 1 and 2 ICEs (pangenomeGraph.gexf) were updated to integrate additional category-level annotations to enable coloring of protein family nodes based on predicted function. Updated gexf files were then imported into the Gephi Lite online tool (https://lite.gephi.org/v1.0.2/) and adjusted as follows. Appearance: Nodes [“Size” set from “nb_genomes”, option selected to “interpolate between custom min and max values” with 10 (min) and 40 (max) selected, otherwise defaults]; Edges [“Size” set from “nb_genes”, option selected to “interpolate between custom min and max values” with 2 (min) and 10 (max) selected, otherwise defaults]. Layout: ForceAtlas2 [“Edge weight influence” 1, “Gravity” 0.05, “Scaling ratio” 10, “Slow down?” 6.5, option for “Strong gravity mode?” selected, otherwise defaults], layout options were defined by first enabling the function to pre-populate with optimal settings for each graph and then setting the Slow down to 6.5 for both. Two versions of each graph were produced with Gephi Lite for each ICE group, one with nodes colored by pangenome partition and one with coloring by functional annotation category. Gephi Lite graphs were then imported into Gephi Desktop v0.11.2 for optimization for display as figures and export as SVG, as follows. In Overview, Appearance settings for Nodes were set by selecting Size and Ranking based on “nb_genomes” from 20 (min) to 100 (max); in Preview, Preview Settings for Nodes were set to select option for “Fixed Border Width”, setting color to black, and for Edges by selecting options to “Show Edges”, “Use weight”, “Rescale weight” from 1 (“Min. rescaled weight”) to 10 (“Max. rescaled weight”), color set to dark grey (#999999) and option for “Curved” unselected.

### Menaquinone pathway analysis

Association of menaquinone pathway genes *menFDBCE* with ICEs in Pg, was evaluated as follows. All *Pg*_set_79 genomes were evaluated for the presence of each gene based on Bakta^49^ annotations, noting that *menC* is annotated as *rspA*. All *menFDBCE* gene sets in each genome were evaluated for co-localization and found to occur as uninterrupted runs of adjacent genes (26 co-localized sets, accounting for all *menFDBCE* genes in *Pg*). Each gene set was then considered in light of available information from curation of ICEs and ICE conjugation module regions. We found that: 8/26 occurred on curated ICEs; 11/26 are within expected regions near annotated ICE conjugation module regions; 4/26 occurred in regions containing canonical ICE genes, however they were not called by ICEfinder^31^ (and therefore excluded from our curated studies), likely due to extensive gaps in the assemblies (three had >100 contigs in original assemblies) or being near or possibly embedded within another ICE; 3/26 occurred on regions that lacked canonical ICE genes, however nucleotide BLAST searches of their concatenated *menFDBCE* genes to *Pg*_ICE_pilot_set_111 revealed matches to curated ICEs with 100% query coverage and >98% identity for each.

### Clinical sample collection

Each potential donor to the study underwent a screening visit, consisting of a full mouth periodontal assessment, determination of eligibility, and collection of pooled subgingival plaque for qPCR screening to determine presence of Pg. Inclusion criteria for enrollment in the study included: age at least 25 years; having at least 16 teeth, excluding third molars; having at least 4 sites with probing pocket depths equal to 5 or 6 mm and concomitant clinical attachment loss; and having detectable Pg in subgingival plaque as determined via qPCR. Enrolled subjects underwent a baseline visit, including a medical and dental history questionnaire and collection of subgingival plaque from the mesial and distal surfaces of posterior teeth. Samples collected from each quadrant were pooled in a sterile cryopreservation solution consisting of 75% medium and 25% glycerol (vol/vol). The medium contained 37 g L-1 of brain heart infusion, 0.004 g L-1 l-cysteine and 0.01% v/v dithiothreitol. Samples were stored at -80°C until used for cultivation. All study procedures were approved by the University at Buffalo Institutional Review Board under protocol STUDY00005287.

### Isolation of strains from clinical samples

To establish the clinical strain collection, an initial set of subgingival plaque samples from eight study subjects from quadrant 1 at an early visit (month 2) were processed. To gain both temporal and spatial insight, subgingival plaque samples from quadrant 3 during a later visit (month 7 or 8) were also processed, under anaerobic conditions, for four of these subjects. Prior to plating, the samples were first homogenized by allowing the material to melt and repeatedly pipetting to disperse visible clumps. The samples were then diluted to 10^-1^, 10^-2^, 10^-3^, and sometimes 10^-4^ their original volume to maximize the opportunity of isolating individual colonies. 100 μL of each sample dilution was then spread on three varieties of BHI blood agar plates [base formula: Brain Heart Infusion (BD Difco Bacto 237500) – 37 g/L, yeast extract (VWR J850) – 5 g/L, L-cysteine (Sigma-Aldrich C7352) – 0.5 g/L, sodium bicarbonate (JT Baker 3506-01) – 1 g/L, and agar (BD Bacto 214010) – 10 g/L; then supplemented (post-autoclaving) with hemin (Sigma-Aldrich 51280) – 1 mL/L (5 mg/mL stock concentration), 1,4-dihydroxy-2-naphthoic acid (TCI D2296) – 10 mL/L (0.1 mg/mL stock concentration), defibrinated horse blood (Hemostat DHB100) – 50 mL/L, and N-acetylmuramic acid (Cayman Chemical 31330) – 1 mL/L (10 mg/mL stock concentration] to optimize the recovery of *Pg*: 1) vancomycin, nalidixic acid, and colistin added to base formula after autoclaving [vancomycin (Sigma-Aldrich V2002) – 1mL/L (2.5 mg/mL stock concentration), nalidixic acid (Sigma-Aldrich N8878) – 1 mL/L (15 mg/mL stock concentration), and colistin (Sigma-Aldrich PHR1605) – 1 mL/L (10 mg/mL stock concentration]; 2) nalidixic acid and colistin added to base formula after autoclaving [nalidixic acid (Sigma-Aldrich N8878) – 1 mL/L (15 mg/mL stock concentration) and colistin (Sigma-Aldrich PHR1605) – 1 mL/L (10 mg/mL stock concentration]; and 3) no antibiotics added to base formula. The spread plates were then incubated anaerobically for 8 days at 37 °C. After incubation, hundreds of colonies were collected for each sample across the three different plate types with a sterile toothpick and re-struck as ∼1.25 cm x 1.25 cm patches on BHI blood agar plates with no antibiotics [base formula]. The patch plates were then incubated anaerobically for 6 days at 37 °C. After incubation, a streak through each patch was collected with a toothpick and frozen in 96-well PCR plates containing a 0.5 mg/mL bovine serum albumin [Spectrum AL135] solution for future PCR and sequencing analyses. The rest of the patch was then collected with a toothpick and frozen in 96-well microtiter plates containing a modified ATCC 2722 broth glycerol solution [glycerol (VWR BDH1172-1LP) – 0.25 L, milliQ water – 0.75 L, Tryptic Soy Broth (BD Bacto 211825) – 30 g/L, yeast extract (VWR J850) – 5 g/L, and L-cysteine (Sigma-Aldrich C7352) – 0.5 g/L; then supplemented (post-autoclaving) with hemin (Sigma-Aldrich 51280) – 1 mL/L (5 mg/mL stock concentration) and 1,4-dihydroxy-2-naphthoic acid (DHNA) (TCI D2296) – 10 mL/L (0.1 mg/mL stock concentration)].

### Metagenome DNA extraction and sequencing

24 total *Pg*-Enrichment Culture Mini-Metagenomes (*Pg*-ECMMs) were prepared and sequenced, with each *Pg*-ECMM metagenome comprising 96 colony patches from a given subject. *Pg*-ECMMs were prepared for 8 subjects across two time points, with each of the 8 subjects represented by at least one *Pg*-ECMM for an early visit timepoint (10 total metagenomes, 1-2 metagenomes per subject), and 4 subjects also represented for a later visit time point (14 total samples, 3-4 metagenomes per subject). See Supplementary Data File 2 for details.

To begin preparation of each *Pg*-ECMM, 96-well plates of the patch culture DNA (in BSA) were thawed at room temperature for 15 min. The plates were then vortexed for 15 sec, rotated on a nutator for 1 min, and each well was pipette-mixed 10 times to ensure resuspension of the material. 20 μL of the resuspended material from each well was transferred to a new 96-well PCR plate and heated in a thermal cycler at 95 °C for 6 min. After heating, 5 μL of material from each well was pooled into a single 1.7 mL Eppendorf tube for each plate (∼480 μL total/tube) to be used for DNA extraction.

Pooled DNA from each of the plates was extracted using the MasterPure Gram Positive DNA kit (MGP04100) which is effective for both Gram-positive and -negative bacteria. In brief, to lyse the bacterial cells, 150 μL of TE Buffer and 1 μL of Ready-Lyse Lysozyme solution was added to each tube of pooled sample and incubated at 37 °C for 18-20 hr. After incubation, 1 μL of Proteinase K (50 g/L stock concentration) was diluted into 150 μL of Gram Positive Lysis Solution. The mixture was then added to each pooled sample. The tubes were incubated at 65-70 °C for 15 min, vortexing every 5 min. The tubes were then slowly cooled by first placing them at 37 °C for 3 min and then on ice for 5 min.

To precipitate the DNA from the lysates of the pooled samples, 175 μL of MPC Protein Precipitation Reagent was added to each tube and vortexed vigorously for 10 sec. The debris was pelleted by centrifugation at 4 °C for 10 min at >10,000 x g (Beckman Coulter X-22R centrifuge with F2402H rotor). The supernatants were removed and added to new 1.7 mL Eppendorf tubes. 1 μL of RNase A (5 g/L stock concentration) was added to the supernatants and incubated at 37 °C for 30 min. After incubation, at least an equal volume of isopropanol, relative to the volume already in the tube, was added to each tube. The tubes were mixed by being gently inverted 30-40 times. DNA was then pelleted by centrifugation at 4 °C for 10 min at >10,000 x g (Beckman Coulter X-22R centrifuge with F2402H rotor). After centrifugation, the supernatants were discarded and the remaining DNA pellets were washed with 500 μL of 70% ethanol. The ethanol was then removed and the tubes were left to dry for 1-2 min. The washed DNA pellets were then resuspended in 25 μL of TE Buffer.

Sample library preparation and shotgun metagenomic sequencing were performed on each DNA extraction by Zymo (Irvine, CA, USA), using the lllumina DNA Library Prep Kit (Illumina, San Diego, CA) with up to 500 ng DNA input following the manufacturer’s protocol and using unique dual-index 10 bp barcodes with Nextera adapters (Illumina, San Diego, CA); samples were sequenced on a NovaSeq X platform with 2 × 151 base pair paired-end reads, to a target depth of at least 10 million reads per sample.

### Metagenome sequence assembly

Quality control of the raw Illumina reads for each metagenomic sample was performed with fastp^110^ v0.24.0 [default parameters] (https://github.com/opengene/fastp). The processed reads for each metagenomic sample were then assembled with MEGAHIT^85^ v1.2.9 [default parameters] (https://github.com/voutcn/MEGAHIT).

### ICE distribution analysis

To evaluate the presence of *Pg*-ICE-like TraG sequences in clinical metagenome assemblies we first annotated all 24 metagenome assemblies using Bakta^49^ v1.9.4 [default parameters]. The bacterial host taxonomic classification for each contig of the metagenomic assemblies was predicted with Kraken2^86^ v2.1.3 [default parameters]. A BLAST database was then generated with all of the predicted proteins from the metagenome assemblies. Proteins predicted from the metagenome assemblies were searched with a representative *Pg* TraG sequence (*Pg*W83 GCA_000007585.1_01362 from the Prokka protein calls in HOMD Genomic RefSeq v11.02) using BLASTp with permissive cut-offs [evalue 1e-10 max_target_seqs 5000 matrix BLOSUM62 gapopen 11 gapextend 1 seg no]. The results were filtered to sequences with ≥99% query coverage and ≥65% sequence identity to the query. This approach yielded 51 total sequences, including 21 predicted to be in *Pg* contigs and the remaining 30 in other species in the Bacteroidales, including *Bacteroides ovatus*, *P. asaccharolytica*, *P. cangingivalis*, *P. somerae*, *P. sp. oral taxon 275*, *Prevotella denticola*, *P. intermedia*, *P. melaninogenica*, *P. nigrescens*, *P. sp. oral taxon 475*, and *Tannerella forsythia*.

To evaluate distributions of *Pg*-ICE-like TraG sequences in other bacterial taxa in the *e*HOMD, we searched the *e*HOMD with the same representative TraG protein sequence as above. The search was carried out on the *e*HOMD webserver using the SequenceServer^111^ 2.0.0 functionality with BLASTP 2.16.0+ [Database: PROKKA::ALL Proteins Annotated from all *e*HOMD Genomes V11.02 (faa) (18701740 sequences, 5869425992 characters))] and permissive cut-offs [“Advanced parameters” selected: evalue 1e-10, max_target_seqs 5000, matrix BLOSUM62, gap-open 11, gap-extend 1, filter F]. Results were filtered to those sequences with ≥99% query coverage and ≥65% sequence identity, allowing capture of full-length TraG genes from all ICE groups characterized in this work as well as sequence-similar genes in other genomes (830 sequences). Hits were then further filtered to include only proteins from oral bacteria (‘oral’ association in the ‘Body Site(s)’ field of the *e*HOMD taxon_table), with the exception of also including 7 *B. ovatus* hits meeting the thresholds, as this species was also detected in our clinical enrichment culture mini-metagenomes from gingival crevice samples, above; together, this yielded 518 total sequences. An additional sequence captured among oral hits with ≥99% query coverage and 64.88% identity was included for use as an outgroup in downstream analyses (GCA_024759565.1_00679 *Phocaeicola vulgatus*).

To assess the phylogenetic relationships among *Pg*-ICE-like TraG sequences and evaluate associations with host taxa and presence in clinical samples, we evaluated as follows. All 570 TraG protein sequences (1 outgroup, 51 clinical sequences, and 518 *e*HOMD sequences including 7 from non-oral *Bacteroides ovatus*) were aligned using muscle v5.1 [defaults] and phylogenetic relationships determined using IQ-Tree^112^ v3.1.1 including ModelFinderPlus to determine the rate substitution model [-m MFP, yielding best-fit model Q.PLANT+I+R5, chosen according to Bayesian Information Criterion (BIC)], and bootstrap to assess branch support [-B 1000 –bnni –alrt 1000]. The tree was visualized using iToL and rooted on the outgroup.

Metadata was mapped as follows. The outermost ring indicates source, which sequences from the *e*HOMD shown in black and those from donor samples shown in neon colors; the middle ring indicates the taxon encoding the TraG, for *e*HOMD sequences this is defined by the provided taxon information and for the clinical samples this is defined by the Kraken2^86^ predicted taxon label; the innermost ring indicates the ICE group for all sequences associated with *Pg* ICE conjugation module regions defined in this study. Hyperlinks to the *e*HOMD JBrowse tracks are available on the interactive web version of the tree and are centered on the TraG gene +/- 3000bp.

